# Spatial distribution of insular cliff vegetation and future scenarios in a climate change perspective

**DOI:** 10.1101/2023.11.09.566357

**Authors:** Miquel Capó, Iván Cortés-Fernández, Joshua Borràs

**Author notes:** Corresponding author: Miquel Capó.

## Abstract

Cliff ecology has been studied for decades, providing good information about its high biodiversity values and their vulnerability to climate change. At the same time, literature shows how insular ecosystems are evolutionarily enhanced to present biodiversity hotspots with high endemicity rates, but they are also severely affected by anthropogenic effects. Together, insular cliff communities combine both biodiversity uniqueness and high vulnerability to global change, but few studies have evaluated these particular ecosystems. In this study, our aim was to provide novel information on the spatial distribution of insular cliff vegetation, taking the Balearic islands as a study case, assessing which environmental and climatic variables contribute to the definition of habitat conditions. Also, we used the predictions found in the WorldClim database to predict how the spatial distribution model vary. The map obtained showed that the mountains and coast were the most suitable habitats, especially the mountain ranges of the island of Mallorca. In particular, the high peaks of Serra de Tramuntana englobe the most fitted conditions for rupicolous species. Regarding the future climatic scenarios, in both the pessimistic and optimistic models, the spatial distribution of cliff vegetation remain intact in the mountain ranges where the habitat suitability was high, but disappear or reduce in other less favorable zones for the period 2021-2040. However, the extension of the distribution will be reduced in mid- and long-term periods, until 2081-2100, when the well-suited areas are only predicted to be in high-elevation mountains from the north of Mallorca island. These results emphasize the high vulnerability of these habitats to climate change due to their vulnerability to aridity and the strict requirements for habitat suitability (*e*.*g*., high slopes). From this, future studies should focus on single-species analysis to evaluate if any cliff specialist species can be at risk of extinction due to climate change.

## INTRODUCTION

Evolution allowed plants to colonize a wide range of habitats around the world. In particular, cliffs are considered as defeating environments where the arrival, establishment, and functionality of plant ecosystems is not favorable due to its abiotic stresses such as excessive irradiation or soil drought among others (Aronne et al., 2018). However, the geomorphological heterogeneity of the cliffs facilitates the presence of microhabitats, which are essential for the development of good plants. For instance, cliffs usually present rock cracks where fertile soil and water are accumulated, and therefore plants present more favorable conditions compared to neighboring areas (Larson et al., 2000). Due to this selective pressure, the composition of cliff plant communities is unique and usually harbors endemic species that cannot be found anywhere else both in geographical and ecological terms (Larson et al., 2000; March-Salas et al., 2023).

All these particular conditions make cliffs a hotspot of biodiversity, including from generalist species to endemic, threatened, or rupicolous specialists (Larson et al., 2000; March-Salas et al., 2018). This diversity make is usually hard to manage in terms of conservation biology, as endemic and threatened species are usually the main target of programs (Gaston, 2010) but generalists are found to be equally affected by threatens such as rock climbing (Lorite et al., 2017; March-Salas et al., 2018).

Cliff plant communities are diverse in terms of functionality, which enhances mutualistic and antagonistic interactions and creates unique ecosystem functioning (Larson et al., 2000). For example, some studies reported the beneficial effect of co-existence on sharing bacterial and fungal symbionts in cliff communities (Krah & March-Salas, 2022) or on facilitating nutrients supply (García-Callejas et al., 2021). In contrast, antagonistic effects have also been reported on cliffs, especially competitive interactions for colonization of suitable microhabitats (do Carmo et al., 2016). These habitats are even more ecologically interesting in areas where evolutionary processes are accelerated, such as the case of islands. There, endemic species rates, population genetics, and biodiversity are substantially higher than mainland regions (Thompson, 2020). Despite all the knowledge about cliff habitats and insular ecosystems, the functioning of insular cliff plant communities remains mainly unknown.

Conservation strategies to palliate global change impacts on the cliff flora are difficult to design due to the scarce information on its ecology (March-Salas et al., 2023) being even more urgent on islands. For example, global change problems are reported in mainland cliff communities, such as colonization of invasive alien flora (Cousins & Witkowski, 2012) or the effect of climate change (Ferreira et al., 2021). In this sense, how global change affects island cliff habitats is urgently needed to optimize and design conservation strategies, especially focusing on endangered and threatened species.

In terms of biodiversity, islands are considered as hotspots (Kreft et al., 2008) and in Mediterranean regions are defined as hotspots inside hotspots (Medail & Quezel, 1997). As a result, they englobe high endemic rates (Guardiola & Sáez, 2023) and act as refuges for many threatened species (Thompson et al., 2005). However, these areas are also affected by anthropic disturbances and global change that comprises the structure and function of insular ecosystems (Leclerc et al., 2020). For example, the impact of alien flora has been recently reported (Cerrato et al., 2023) as well as the effect of climate change (Cubas et al., 2022; M. T. Ferreira et al., 2016). Other indirect effects on ecosystem functioning have also been reported, such as pollinator or seed disperser loss on islands (Traveset & Navarro, 2018) or the alien herbivores impacts (Capó et al., 2023; Cubas et al., 2019).

Due to the lack of information on the ecology of cliff vegetation on islands, we took the opportunity to study the ecology of the rupicolous flora of the Balearic Islands. This archipelago is located in the western Mediterranean Basin, close to the eastern Iberian coast. It consists of five islands: Mallorca (3640 km^2^), Menorca (701 km^2^), Eivissa (541 km^2^), Formentera (82 km^2^) and Cabrera (13 km^2^) together with ca. 150 islets (Guardiola & Sáez, 2023). The archipelago is a good example of continental Mediterranean islands with a high rate of endemic species (approximately 6.9 – 10.4%) (Guardiola & Sáez, 2023; Sáez et al., 2011) and it is also affected by global change impacts (Abdallah et al., 2021; Capó et al., 2021; Ramis et al., 2017; Traveset et al., 2008). However, the ecology of its rupicolous flora is mostly unknown (Tomas et al., 2019) despite the floristic inventories indicating a high proportion of endemic flora that are only found in rupicolous habitats (Llorens et al., 2007; Rita & Payeras, 2006; Sáez et al., 2017). Additionally, these areas present a high number of threatened species, some of which are considered extremely narrow (Fenu et al., 2020; Guardiola & Sáez, 2023; Sáez & Roselló, 2004). In this context, our aim was to (1) determine which abiotic variables characterize the geographical distribution of rupicolous in the archipelago; (2) define the spatial hotspots of rupicolous flora in the archipelago; (3) predict the geographic refuges for insular cliff species in a climate change scenario. We expect that the most potential areas for rupicolous flora will locate in Mallorca due to its heterogeneous topography and orography, whereas on the island we expect that areas with moderate temperatures and precipitation regimes will maximize the probability of occurrences rather than extreme values of both parameters. Taking into consideration the climatic change scenario, we expect that potential habitats will be scarcer and only some particular sites will act as climatic refuges.

## MATERIAL AND METHODS

### Study area

The Balearic Archipelago consists of five islands and about 150 islets (see above), which are englobed in the western Mediterranean region. Mallorca is the largest island and contains three geomorphological spots: Serra de Tramuntana and Serra de Llevant, consisting of the main mountain ranges; and the central lowland (Guardiola & Sáez, 2023). Mountain ranges are not as tall as the analogues of the mainland, but some peaks of Serra de Tramuntana are higher than 1000 m and the highest peak is 1445 m. In the case of the other islands, the mountain ranges are scarce and only some eventual mountains reach considerable altitudes (the highest peak of Menorca is 358 m, Eivissa 475 m, and Formentera 202 m). The orography of all islands allows the presence of rupicolous communities, included as a priority habitat (8210 Calcareous rocky slopes with chasmophytic vegetation). These habitats are characterized by the importance of precipitations and temperature to the creation and maintenance of suitable areas where plant communities can be stablished (Fornós et al., 2009)

The climatology is mainly Mediterranean, where precipitations are concentrated during spring and autumn, and temperatures reach the highest values in summer. Precipitations substantially vary between islands and elevations: maximum precipitations are recorded in Serra de Tramuntana (1500 mm / year), while minimum precipitations are recorded in coastal areas from Eivissa and Formentera (less than 300 mm/year) (López Mayol et al., 2017; Ramis et al., 2017). In fact, Serra de Tramuntana has been considered the wetter areas of the Balearic Islands (humid and subhumid ombrotypes), while southern Mallorca, Eivissa, and Formentera are considered dry and semiarid ombrotypes (Llorens et al., 2007).

### Data Collection

The specialization of the rupicolous depends on each species, as while some of them can live on cliffs but are mainly located in other habitats, others are cliff specialists (March-Salas et al., 2018). Hence, to properly characterize cliff habitats, we selected 20 species that are considered cliff specialists on the general literature (Castroviejo, 1986) with various geographical distribution ranges, accounting for a total of 242 occurrences (Table 1).

**Table 1.** Selected species for ENM analysis.

| Species | Family | Distribution | No. occurrences |
| --- | --- | --- | --- |
| *Arenaria grandiflora* ssp. *bolosii* | Caryophyllaceae | Endemic | 4 |
| *Brassica balearica* | Brassicaceae | Endemic | 14 |
| *Bupleurum barceloi* | Apiaceae | Endemic | 1 |
| *Cephalaria squamiflora* ssp. *balearica* | Caprifoliaceae | Endemic | 6 |
| *Cheirolophus intybaceus* | Asteraceae | Widespread | 17 |
| *Dianthus rupicola* ssp. *bocchoriana* | Caryophyllaceae | Endemic | 1 |
| *Genista majorica* | Fabaceae | Endemic | 10 |
| *Helichrysum crassifolium* | Asteraceae | Endemic | 29 |
| *Hippocrepis balearica* | Fabaceae | Endemic | 67 |
| *Laserpitium gallicum* ssp. *majoricum* | Apiaceae | Endemic | 1 |
| *Lomelosia cretica* | Caprifoliaceae | Widespread | 50 |
| *Lonicera pyrenaica* ssp. *majoricensis* | Caprifoliaceae | Endemic | 3 |
| *Pimpinella tragium* ssp. *lithophila* | Rosaceae | Widespread | 2 |
| *Potentilla caulescens* | Rosaceae | Widespread | 14 |
| *Primula vulgaris* ssp. *balearica* | Primulaceae | Endemic | 3 |
| *Sesleria insularis* | Poaceae | Endemic | 8 |
| *Silene mollissima* | Caryophyllaceae | Endemic | 3 |
| *Teucrium cossonii* | Lamiaceae | Endemic | 4 |
| *Thymus richardii* | Lamiaceae | Endemic | 1 |
| *Viola jaubertiana* | Violaceae | Endemic | 4 |

Data obtained from our own knowledge were complemented with species occurrences obtained from GBIF database (The Global Biodiversity Information Facility, 2023) and from the public biodiversity database BiodiBal of the University of the Balearic Islands (Biodibal, 2023). After data acquisition, a rigorous selection of good quality data was performed. We selected only living specimens occurrences and excluded data with >10 m uncertainty prior to 2010s. Furthermore, we used all points from the Poaceae family from GBIF, with the same quality selection, as a proxy for oversampling of some area (*e*.*g*., Mallorca vs. other islands) (Figure 1). We selected this family since most are only reported by an expert botanist or exhaustive census of certain areas.

**Figure 1.**
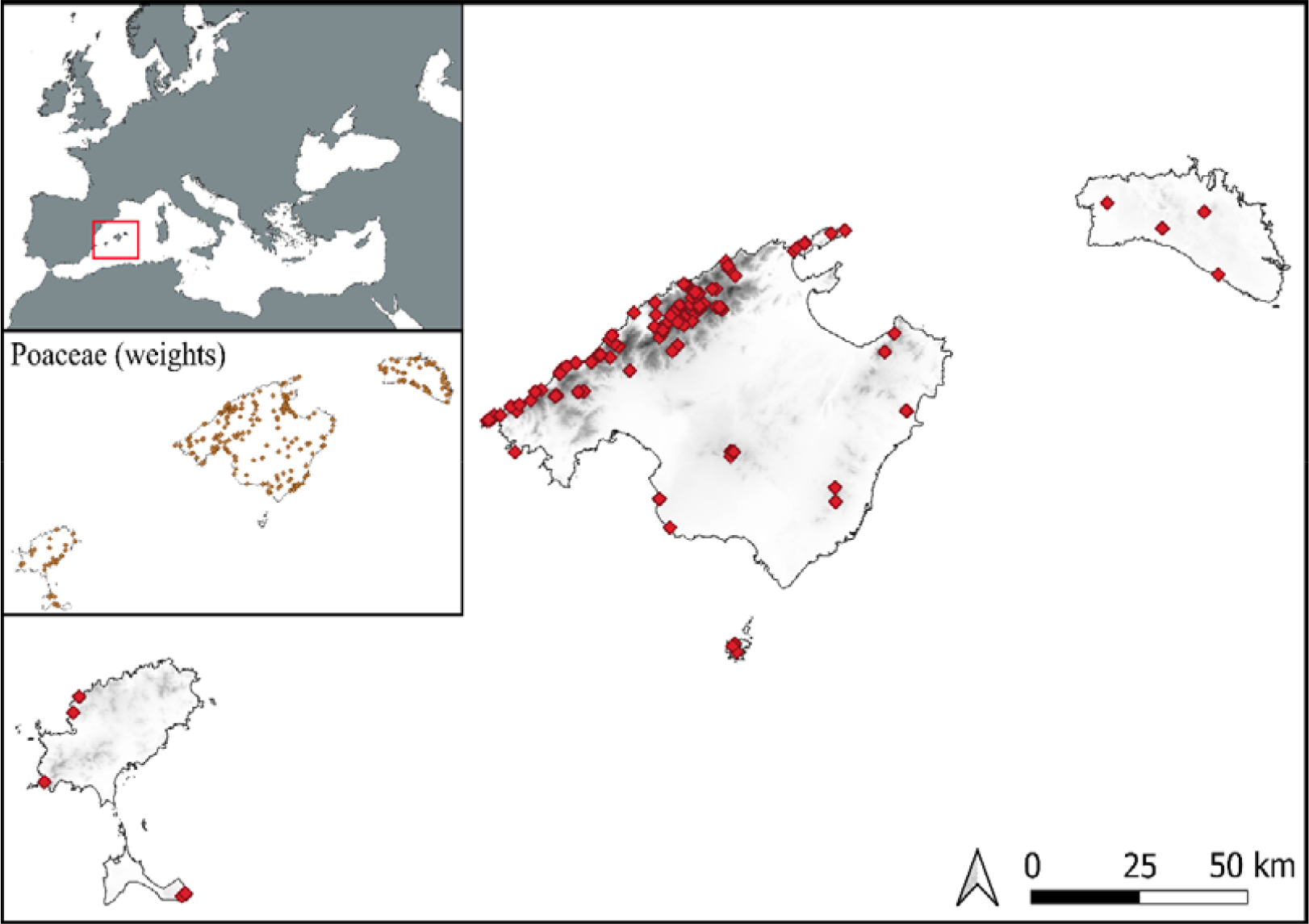
Balearic Islands archipelago indicating plant occurrences (n = 242) collected from GBIF, BiodiBal and personal knowledge. The elevation is displayed in gray scale.

### Enviromental variables

Based on our ecological knowledge of the rupicolous communities, 26 environmental variables were selected (Table 2). The 19 bioclimatic variables were obtained from WorldClim 2.1 (Fick & Hijmans, 2017). Elevation, slope, distance to shoreline, and orientation NS and EW were calculated using ‘*r*.*slope*.*aspect*’ function from GRASS (GRASS Development Foundation, 2022) with the 5 m Digital Terrain Models (DTM) a LiDAR-based product, for distance to shoreline we also used the shoreline shapefile and land use information from SIOSE AR. All these files are obtained from the Spanish National Geographic Institute (Instituto Geográfico Nacional, 2021). Orientation calculated from DTM05 was transformed to NS and EW using sin and cos respectively (Grenon & Laflamme, 2011). The geology of the surface materials (hereafter, geology) were obtained from GEODE offered by the Spanish Geological Survey (GEODE, 2023). All the resulting layers had identical coordinate systems and a spatial resolution of 30 arcsec spatial resolution (∼1 km^2^).

**Table 2.**
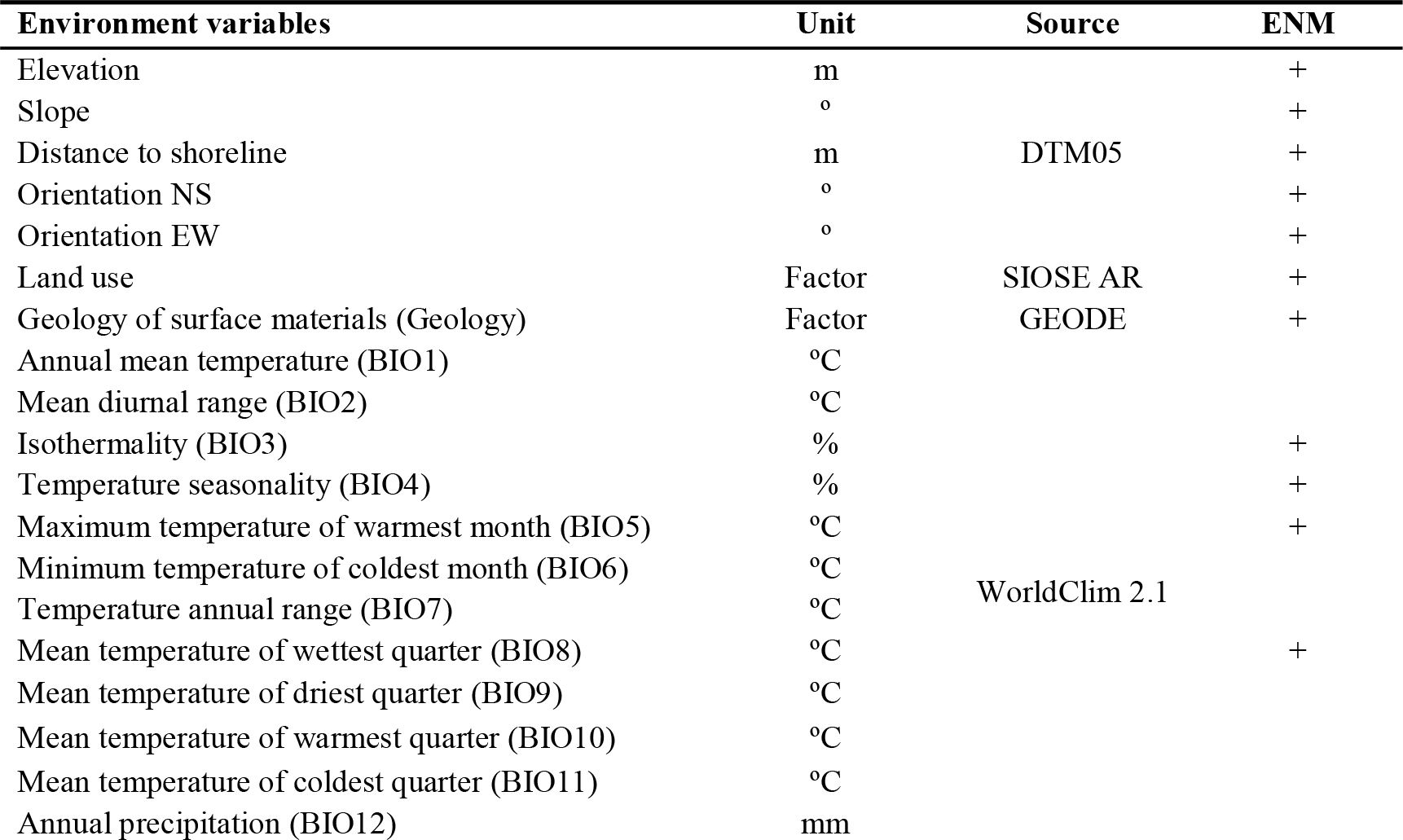
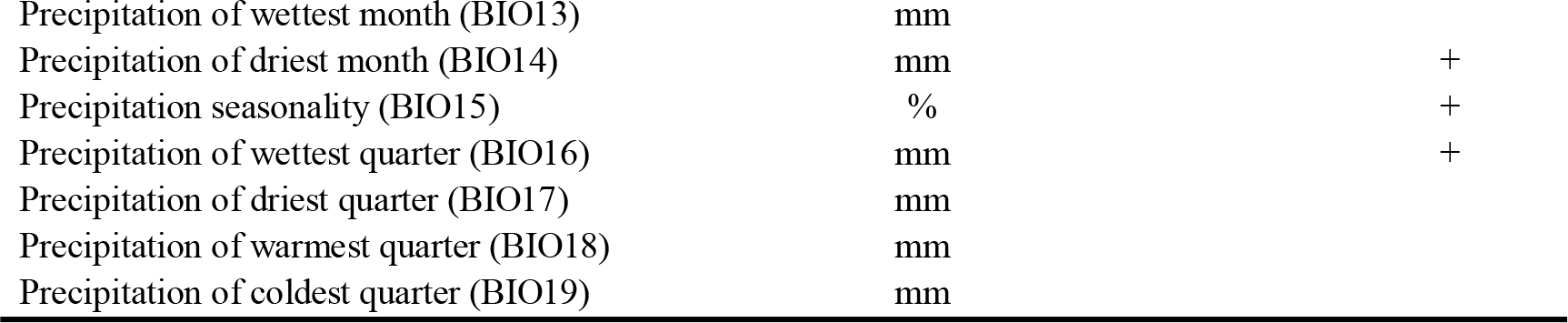
Environmental variables used in modeling. Variables used in ENM analyzes are marked with ‘+’.

To avoid multicollinearity of the 26 environmental variables, we applied a pre-selecting variables approach within each set of environmental variables. We eliminated, one by one, the predictor variables that yielded the highest variance inflation factor (VIF) values until all variables had a VIF below 50 (Dormann et al., 2013). This approach allowed us to reduce the risk of multicollinearity, but maintain significant ecological variables. Among the 26 original environmental variables, only 14 variables were ultimately selected for modeling, as reported in Table 2.

For future climatic predictions, we used the bioclimatic variables of WorldClim for the four-time intervals (2021-2040, 2041-2060, 2061-2080 and 2081-2100) based on four projections for four Shared Socio-economic Pathways (SSPs): 1-2.6, 2-4.5, 3-7.0 and 5-8.5 (Fick & Hijmans, 2017). Two different simulations of the future climate developed by the Euro-Mediterranean Center on Climate Change (hereafter, CMMC)(Lovato et al., 2022) and Max Planck Institute (hereafter, MPI)(von Storch et al., 2017) were used. These projections were chosen because they were the worst case (CMCC) scenario and the best case (MPI) scenario within the area studied.

### Random forest modeling

Data manipulation and statistical analyzes were performed using the R version 4.3.1 statistical software version 4.3.1 (R Core Team, 2023). We used a classification algorithm to model the species distribution because our study focused more on the discrimination capacity of ecological niche models (hereafter ENM). Random forest (hereafter, RF) models need both presence and absence data, but we lacked absence data in this study. Hence, we generated double the number of background points (n = 484), using the ‘*spatSample’* function from ‘*terra’* R package (Hijmans, 2023). We assumed that any points located at least 1 km from presence data could be considered as pseudo-absence data.

Rupicolous communities can be found on coastal cliffs, therefore very close to the coastline. Due to resolution issues, some of these records may fall into no-data pixels of the environmental set of predictors, that is, they are sea pixels in the predictors’ layers. For each predictor variable, we assigned the value of their closest neighbor to these no-data pixels using the *‘preProcess’* function from *‘caret’* R package (Kuhn, 2008). This allowed us to keep all occurrence records for modelling, instead of having to discard valid occurrence points from already scarce data.

We used 40-fold cross-validation combined with 40 replications to build RF models with *‘ranger’* implementation through the *‘train’* function in ‘*caret’* R package (Wright & Ziegler, 2017). The number of trees was set to 500, while the number of variables tried at each split, and the minimum size of terminal nodes was obtained by a pre-training performed with *‘tuneRanger’* from the *‘tuneRanger’* R package (Probst et al., 2019). The resulting averaged evaluation of the RF model (AUC and Kappa) was already reported by the *‘train’* function. The probability threshold of the RF model was calculated with the *‘thresholder’* function of the ‘caret’ R package. We selected the minimum threshold with precision value above 95%. For current and future ENM, we constructed predictor layers with all environmental variables only changing bioclimatic variables. ENM prediction layers were generated with the *‘predict’* function from *‘terra’* package.

## RESULTS

### Current species distribution model

The resulting RF model had a prediction accuracy with a mean (± SE) AUC value of 0.9419 (± 0.0557), a mean (± SE) kappa coefficient of 0.8684 (± 0.1265) and an OOB error of 4.76%. We calculated the threshold of significant probability of presence (0.45), as the minimum threshold with a precision value greater than 95%. The predicted ENM with our RF model had a positive accuracy of 95.25% and negative accuracy of 90.84%. The predicted ENM had an extension of 385.92 km^2^, with a mean (± SE) probability of presence of 0.6909 (± 0.0016) .

The appearances of the 20 rupicolous species were mainly distributed in the mountains of Mallorca. The occurrences were mostly located along the Serra de Tramuntana in the northern part of Mallorca. In particular, Puig Major (highest peak) and surrounding mountains located at the heart of the Serra de Tramuntana showed the highest value scores. Meanwhile, many occurrences were found in the Serra de Llevant mountain range on the north-east side of Mallorca, isolated mountains from the center of the island, such as Puig de Sant Salvador, Puig de Randa, Santueri (Mallorca), and in mountain spots of the other islands (Figure 2). The rupicolous spots were found not only in mountains but also in coastal cliffs along the archipelago. As expected, no occurrences were found in the lowlands of the archipelago.

**Figure 2.**
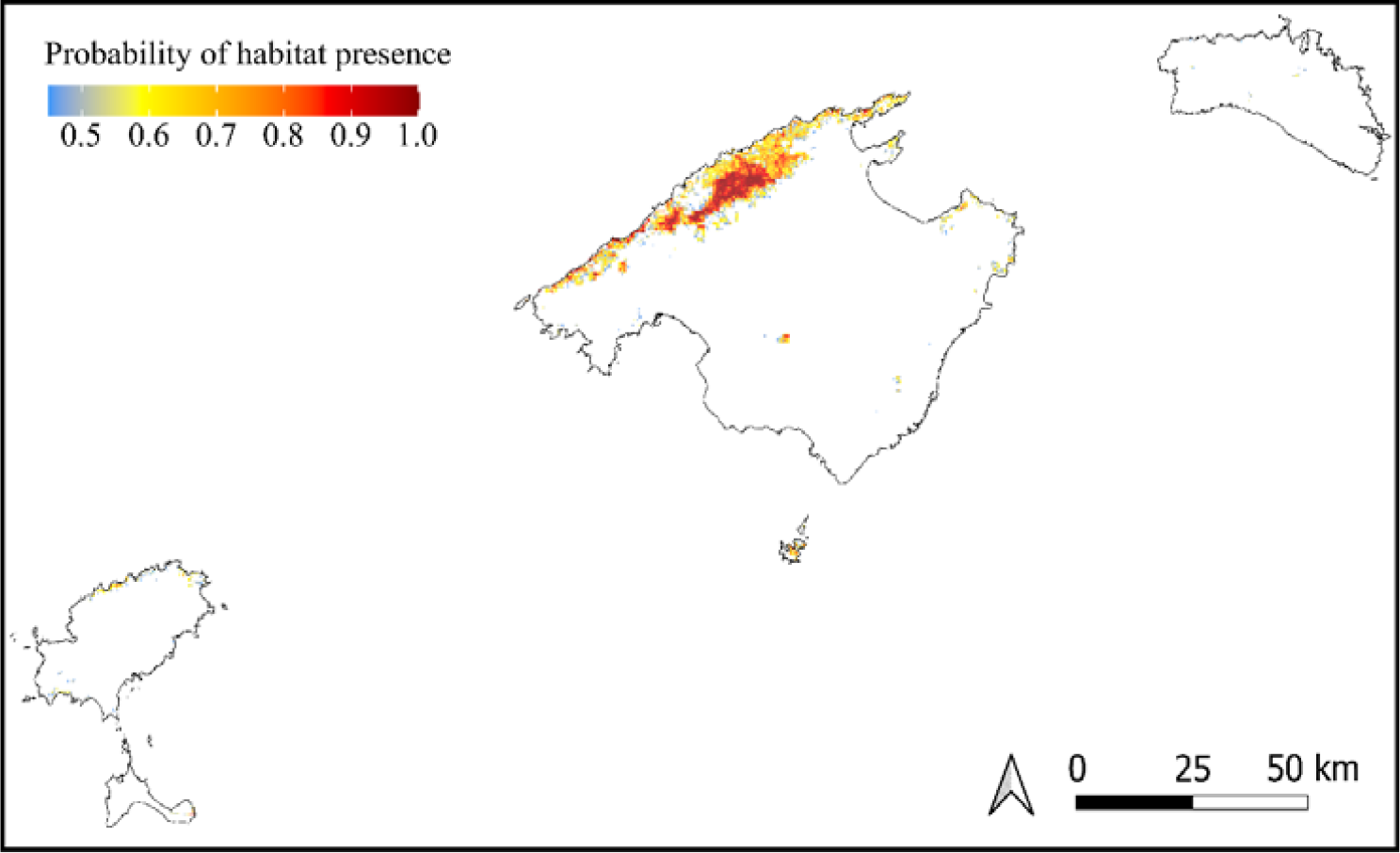
Distribution map of the rupicolous communities from the RF model. Only the probabilities above the minimum threshold with a precision of 95% (0.45) are shown. The positive accuracy of the prediction was 95.25% and the negative accuracy was 90.84%. The model was calculated with 242 occurrences and 484 pseudo-absences.

The importance and response curve of the environmental variables are shown in Figure 3 (and Supplementary Figure 1). Of all environmental variables used to generate the ENM, the slope was the most important variable (Gini = 64.88), however, all climatic variables related to the mean temperature of the warmest month (BIO5), *i*.*e*., August (Gini = 39.06) and the wettest quarter (BIO8), *i*.*e*., autumn (Gini = 33.84) followed in importance in the resulting RF model. The response curve shows that rupicolous communities can be found at the bottom of cliffs (0% slope), and mostly at slopes higher than 15%. Meanwhile, the mean temperature of both the warmest month and the wettest quarter showed higher predictions at the lowest range of temperature. The response curve for the distance to shore showed that most communities are found in coastal cliffs, with a second increase corresponding to the Serra of Tramuntana distance to shore, also shown in the elevation response curve. Finally, the response curve for precipitation seasonality (BIO15) showed to a maximum of less than 44% and over 52%. These two maxima coincide with the two types of zone that the rupicolous communities inhabit. The first, Serra de Tramuntana, has a seasonality of precipitation in the lower part of the spectrum (∼ 40%). Meanwhile, rupicolous communities in smaller islands (*i*.*e*., Cabrera and Formentera), and the coastal cliff in the south of Mallorca, have values at the upper end of the spectrum (∼ 54%).

**Figure 3.**
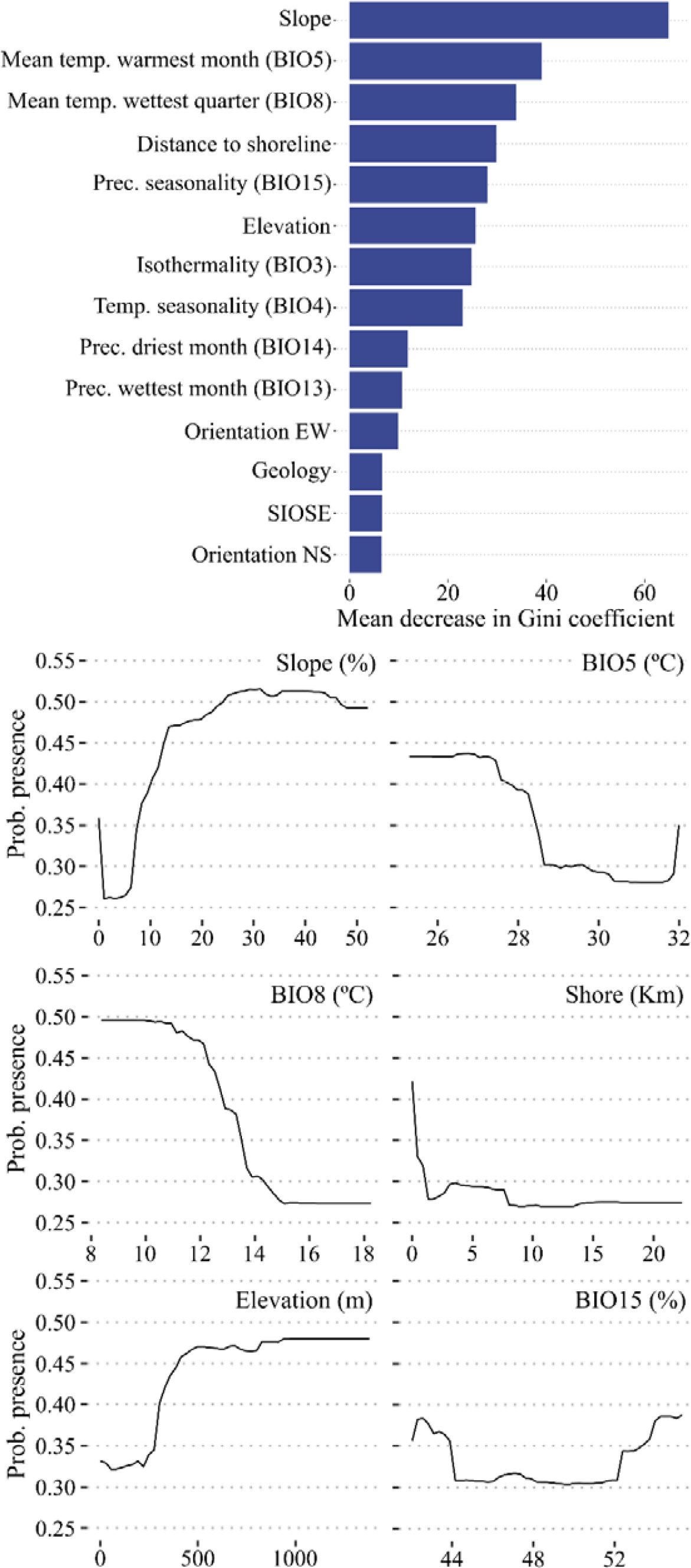
The importance of environmental variables that affects the occurrence of rupicolous communities derived from the RF model. Response curves for the RF model are shown for the top 6 variables; all response curves are shown in Supplementary Figure 1.

### Future species distribution models

All future predictions showed a reduction in the potential habitat of rupicolous communities (Figure 4 and Supplementary Figures 2 and 3). All predictions showed that the potential habitat of all islands and islets except for Mallorca will be greatly reduced. Although some predictions may show an increase in habitat extension, the probability of presence for all predictions decreased compared to the current probability ENM. Both the worst-case (CMCC) and best-case (MPI) scenarios showed a reduction in the mean (± SE) probability value with a range from 0.6073 (± 0.0011 in SSP2-4.5 at 2021-2040) to 0.4973 (± 0.0004 in SSP5-8.5 at 2081-2100); and from 0.5902 (± 0.0010 in SSP3-7.0 at 2021-2040) to 0.5212 (± 0.0007 in SSP5-8.5 at 2081-2100); respectively.

**Figure 4.**
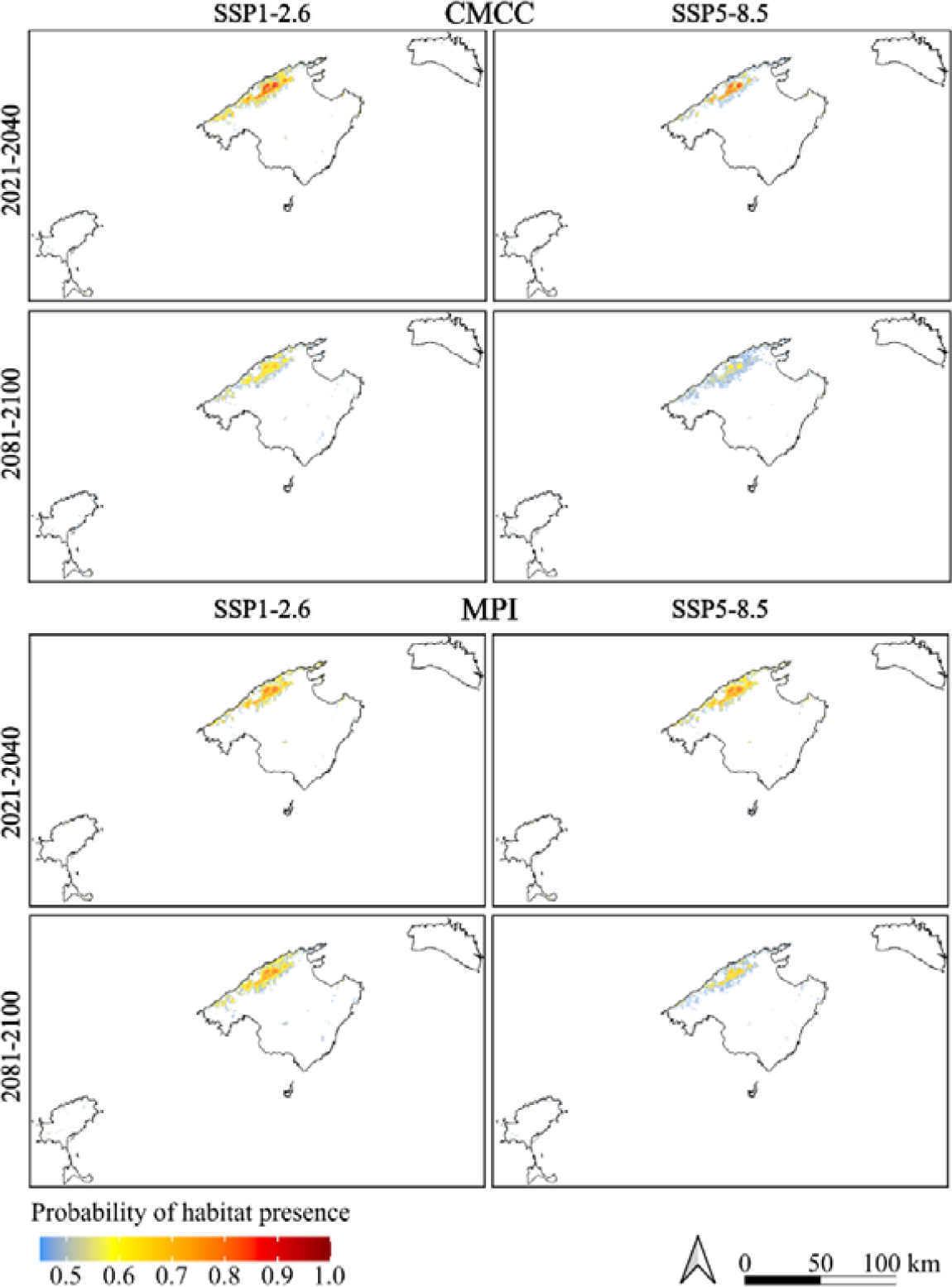
Future distribution prediction of the rupicolous communities from the RF model. The prediction of CMCC and MPI is shown as a worst- and best-case scenario, respectively. All the predictions for the four SSP and four two-decade intervals can be found in Supplementary Figures 2 and 3.

## DISCUSSION

### Spatial distribution of insular cliff vegetation

This study has provided the first ENM of cliff vegetation in the Balearic archipelago, and, to our knowledge, the first of insular ecosystems of cliffs. This information allows us to understand which areas are more suitable for cliff vegetation, as well as their environmental and climatic requirements. This assessment allowed us to predict how these communities would vary under different scenarios of climate change and to detect the most probable climatic refuges.

The current spatial distribution of cliff vegetation is mainly found in mountain ranges and coastal cliffs, due to the presence of suitable habitats for rupicolous vegetation (Larson et al., 2000; Strumia et al., 2020), which are not found in lowlands or small islets. Interestingly, the distribution of cliff vegetation also includes isolated mountains with an altitude of about. 500 m. Indeed, the rupicolous flora can also be found on small islands such as Dragonera or Cabrera, where suitable habitats are also present, despite their low elevation. This is common in rupicolous plant communities, as specialization is not always linked to elevation (Cooper, 1997), but it depends on the study species particular ecology (Aronne et al., 2015; Jung et al., 2019).

In our model, the most important variable was the slope, the mean temperature of the warmest month (BIO5), and the mean temperature of the wettest quarter of the year (BIO8). This validates the strict topographic conditions needed by rupicolous communities and their vulnerability to temperature variation (Larson et al., 2000). Indeed, aridity and drought have been shown to be the main limitation for the survival and growth of cliff species in other studies, such as the case of *Primula palinuri* (Aronne et al., 2018) and even generalist cliff species from the Mediterranean coasts (Ciccarelli et al., 2016).

Other ecological factors can also influence the level of specialization of rupicolous communities, causing an underestimation of their distributional potential. For example, the overpopulation of introduced goats in the mountain ranges of Mallorca has forced many endemic species to be rupicolous to escape from predation (Capó et al., 2023) and some species might be distributed in other habitats in the absence of herbivores. However, there is no information on the distribution of these species prior to the high herbivore pressure in these zones, which limits our analysis to the current situation. Further studies should assess how the island’s rupicolous species can reduce their level of specialization if the population of introduced herbivores is reduced or eradicated. Also, orientation was not important for ENM construction, but some species are unique to the north sides of mountains (*e*.*g. Hippocrepis balearica*) while others are found on the south side (*Lomelosia cretica*) emphasizing the importance of considering species-specific characteristics to assess particular cases (García et al., 2020; Marcer et al., 2013). In fact, this study acts as a first general step in the future to analyze species-specific ecological niche models.

### Future scenarios of cliff vegetation spatial distribution

The projected ENM in both scenarios (MPI and CMCC) predicts a substantial decrease in habitat suitability for cliff vegetation. As observed in other studies, climate refuges are located in high-altitude areas such as the peaks of mountain ranges (Randin et al., 2009) in all possible ENM predictions, following the escalator to extinction rule (Urban, 2018). The importance of temperature (BIO5, BIO8, BIO3, BIO4) and precipitation (BIO15, BIO14, BIO13) for the model implementation are responsible for the reduction or disappearance of potential habitat in the driest areas of the archipelago, and only Serra de Tramuntana is expected to be able to host cliff plant communities in the future, as occurs in many other rupicolous species worldwide (March-Salas et al., 2023).

Focusing on the evolution over time, the whole Serra de Tramuntana is predicted to maintain the same habitat suitability for the period 2021-2040, with a slight habitat loss in the lowest locations while SSP scenarios are getting worse. Habitat suitability ameliorates along with temporal series, and the worst prediction is found at the 2081-2100 period in the SSP5-8.5 prediction. This is not surprising, as the main climatic variables of the ENM (temperature and precipitation) will get worse due to the increase in atmospheric CO_2_ (increase in temperature and reduction in precipitation) in all possible scenarios of climate change. This prediction is common in other ENM studies focused on plants (Williams et al., 2009) and even animals (Cerman et al., 2022). Indeed, Guardiola & Sáez (2023) found a similar pattern for all endemic vegetation of the Balearic archipelago.

The use of ENM must be understood as a predictor, but the information cannot be directly extrapolated to one particular species of our dataset. For example, the microclimatic conditions of particular mountains might offer specific well-suited areas for the survival of some concrete species. For example, the narrow-distributed *Dianthus rupicola* ssp. *bocchoriana* is only found in the Formentor peninsula (north-eastern side of Mallorca) and its whole distribution range is predicted to disappear according to the predictions of our ENM. However, this result cannot be understood as an extinction scenario, as other local environmental variables should be considered for this assessment (García et al., 2020).

### In situ and ex situ management applications

The fragile conditions of rupicolous communities have motivated managers to develop conservation programs (Brampton, 1998). The use of ENM contributes to define which are the most important areas to conserve or manage. For instance, Mendoza-Fernández et al. (2022) found that populations found in low elevations were threatened and required conservation strategies when ecological niche models were adjusted at high elevations. In this sense, short-term measures must be taken in low-elevation areas rather than mountain summits, where habitat is supposed to be adequate in the future. These conclusions can also be extrapolated to our findings. Populations of species of rupicolous found in areas away from Serra de Tramuntana or present in other islands must be considered in short-term conservation programs, especially those where other anthropogenic impacts are also occurring (*e*.*g*., introduced herbivores or habitat loss).

Focusing on the long term, while species populations are being forced to colonize high elevations (*i*.*e*., low-elevation populations disappear), their colonization capacity might fail if previous established vegetation are good competitors (Crepaz et al., 2021). Direct studies are necessary to unravel the colonization capacity of high-elevation areas, or if migration assistance will be necessary (Torres et al., 2023).

## CONCLUSIONS

The study obtained an ENM with general trends in habitat suitability of cliff vegetation. The communities are distributed mainly on the mountain and coastal cliffs of the archipelago, but occurrences are mainly located on the island of Mallorca. Of all geographical areas where habitat is suitable for cliff vegetation, the Serra de Tramuntana is the most suitable and acts as a hotspot for populations. In a climate change scenario, the potential habitat is reduced to the mountains of Serra Tramuntana, where favorable climatic conditions would still be present. However, these patterns should not be extrapolated to conservation at the species level, as other local-scale variables play a key role on their survival.

## ACKNOWLEDGEMENTS

The authors are grateful to all naturalists who contributed through citizen science by uploading plant geolocations on open databases. Dr. Carles Cardona and Gabriel Bibiloni provided us with inspiration to understand the Balearic cliff ecology and Dr. Daniel Argüeso for his inspiration and advises during the ENM analysis. MC was supported by the Juan de la Cierva Formación 2021 postdoctoral fellowship (FJC2021 046888 I) funded by the Spanish Ministry of Science and Innovation and the European Social Fund (NextGenerationEU/PRTR). JB was supported by a PhD fellowship FPI/055/2021 through the Direcció General de Política Universitària i Recerca (Govern de les Illes Balears) and the European Social Fund.

## AUTHORS CONTRIBUTION

Miquel Capó: conceptualization (equal); investigation (equal); methodology (equal); writing – original draft preparation (lead); writing – review & editing (equal). Ivan Cortés-Fernández: data curation (equal); formal analysis (equal); investigation (equal); visualization (equal); writing – review & editing (equal). Joshua Borràs: data curation (equal); formal analysis (equal); investigation (equal); visualization (equal); writing – review & editing (equal).

