## Supplementary figures and images for "Spatial distribution of insular cliff vegetation and future scenarios in a climate change perspective"

### Supplementary Figure 1

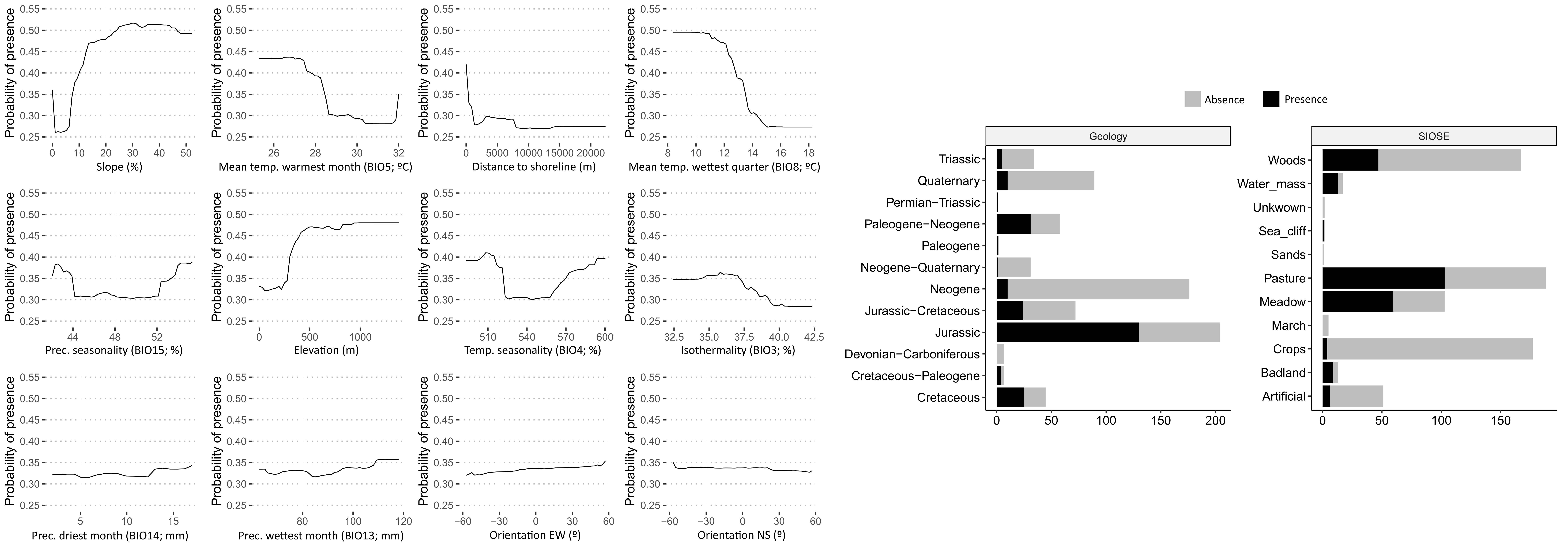

### Supplementary Figure 2

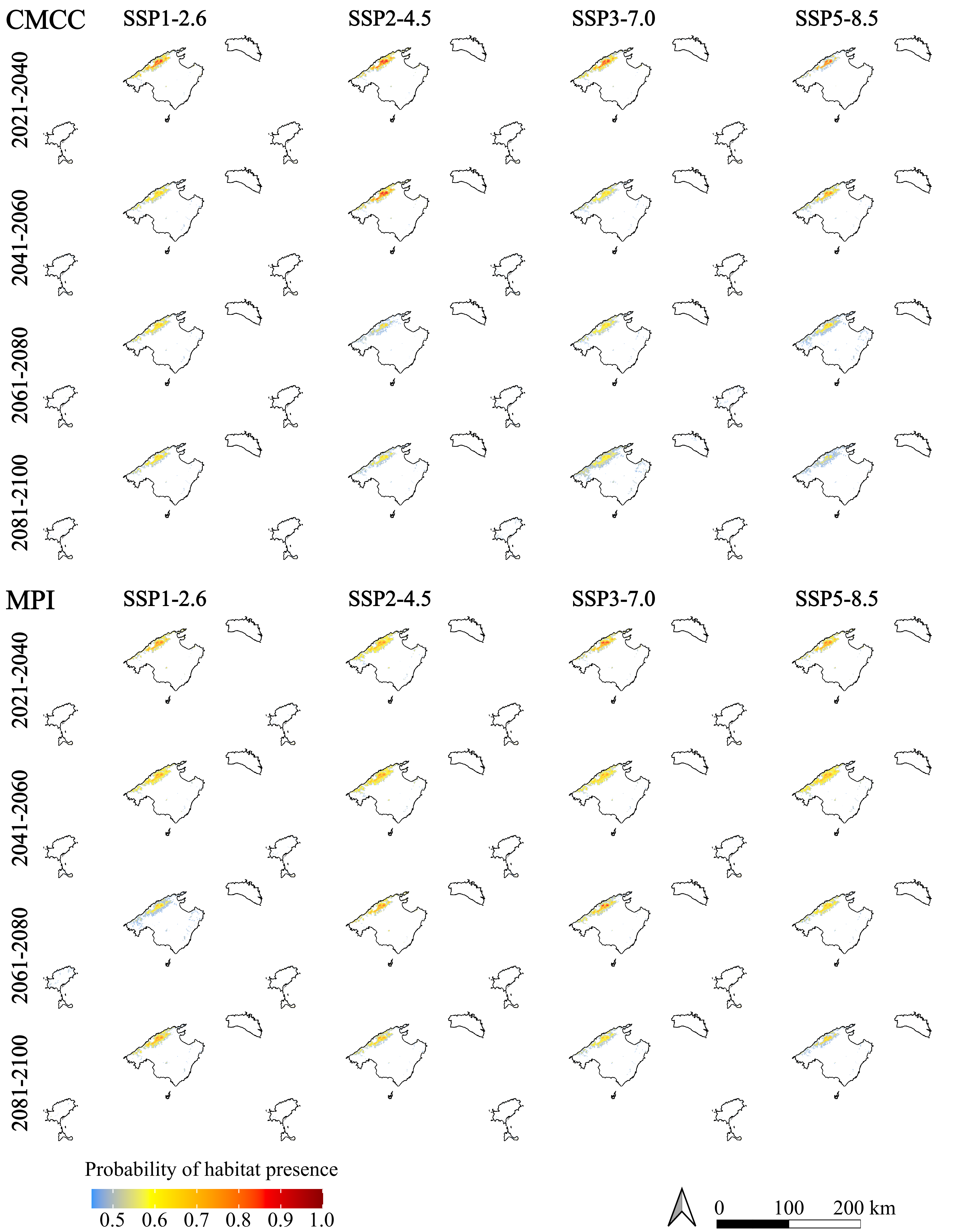
